# Microfluidic-enabled production of DNA barcoded APC library (MEDAL) for high throughput T cell epitope screening

**DOI:** 10.1101/2024.05.07.593072

**Authors:** Xu Cui, Yi Liu, Lih Feng Cheow

## Abstract

Screening for peptide fragments that can be displayed on antigen-presenting cells is an essential step in vaccine development. The current approach for this process is slow and costly as it involves separately pulsing cells with chemically synthesized peptides. We present Microfluidic-Enabled production of DNA-barcoded APC Library (MEDAL), a high throughput microfluidic droplet platform for parallel production of DNA-barcoded APCs loaded with enzymatically synthesized peptides. Droplets containing peptides and their encoding DNA are produced from microfluidic PCR-IVTT reaction. APCs presenting both peptides and DNA barcodes are obtained by injecting cells into these droplets. Up to 9,000 different APCs can be produced and screened within a 10-hour workflow. This approach allowed us to identify peptide sequences that bind to APCs expressing H-2Kb MHC class 1 molecule with next-generation sequencing of DNA barcodes.

## Introduction

T cells are essential players in the adaptive immune system involved in cell-mediated immunity. In our body, T cells can recognize virus-infected or cancer cells by binding to MHC-peptide complexes that are displayed on target cells, and swiftly activate their effector functions such as cytolysis and cytokine secretion to eliminate them. As such, the determination of immunogenic T-cell epitopes is crucial to developing preventive measures such as vaccines and therapeutic interventions such as adoptive-cell immunotherapies (1).

Traditional T-cell epitope mapping efforts based on peptide-MHC multimers and cellular expression (2) are limited in their ability to screen large libraries of peptide-MHC epitopes. Although there have been recent efforts to enhance the multiplexing capacity using fluorescent-(3), mass-(4), and DNA-tagged peptide-MHC complexes (5), efficient production of a diverse set of peptide remains a critical bottleneck to scaling up these assays. Synthesizing large libraries of peptides by chemical methods is expensive and time-consuming (6). In TetTCR-seq (7), peptide synthesis for peptide-MHC multimer production is simplified by performing in-vitro-transcription-translation (IVTT) from DNA templates. However, the requirement to individually synthesize each DNA template, coupled with the large reagent volumes required for IVTT of each peptide and separate peptide purification steps still presents a large barrier to further scaling up the assay.

Microfluidic technologies have found many important applications in immunological screening efforts, due to their ease of miniaturization and automation. In particular, droplet-based microfluidic platforms have been used to screen for immune cells secreting antibodies that bind to specific targets (8) and even neutralize specific viruses (9). Nonetheless, screening for T-cell epitopes is much more complex as it requires the assembly of MHC-peptide complexes, and has not been demonstrated to date.

In this manuscript, we present Microfluidic-Enabled production of DNA-barcoded APC Library (MEDAL), a novel microfluidics platform that would overcome the barriers of high-throughput peptide-MHC complex screening. MEDAL involves first preparing a pool of DNA encoding the peptide of interest. The cost of synthesizing pooled DNA is considerably cheaper than individual DNA or peptide synthesis. Massively parallel clonal amplification of individual DNA templates in the pool is performed via droplet PCR. The high cost of chemical peptide synthesis is replaced by in-droplet enzymatic peptide synthesis from its encoding DNA through the in vitro transcription-translation (IVTT) process. Finally, an antigen-presenting cell (APC) library is prepared by injecting cells into droplets to simultaneously label them with peptides and their encoding DNA. With this integrated microfluidic workflow, a high-diversity DNA barcoded APC library can be prepared in less than 10 h. Through next-generation sequencing of DNA barcodes, we demonstrate the identification of high-affinity peptide sequences that bind to APCs expressing the H-2Kb MHC class 1 molecule.

## Results

Unlike other screening approaches that use peptide-MHC multimers, MEDAL generates a library of APC simultaneously bearing clonal peptide-MHC complexes and DNA barcodes for identification. Compared to synthetic peptide-MHC multimers, APCs can present antigens in more physiological conformation at higher densities (10).

While APCs with different peptide-MHC complexes could be prepared by individually pulsing cells with the corresponding peptides, this approach required individual peptide synthesis and large amounts of reagents for each reaction and would be very costly for large library screening (Fig 1A). We propose to overcome this issue by partitioning a low-cost pooled DNA library into its individual components by droplet PCR, enzymatically synthesizing clonal peptides in each droplet, and loading APCs with these clonal peptides in each droplet (Fig 1B). Using this approach, we expect a significant reduction in reagent cost (∼100 μL in plates vs ∼1 nL in droplets) as well as considerable cost savings in DNA synthesis (pooled vs individual templates) for screening large libraries.

**Fig. 1.**
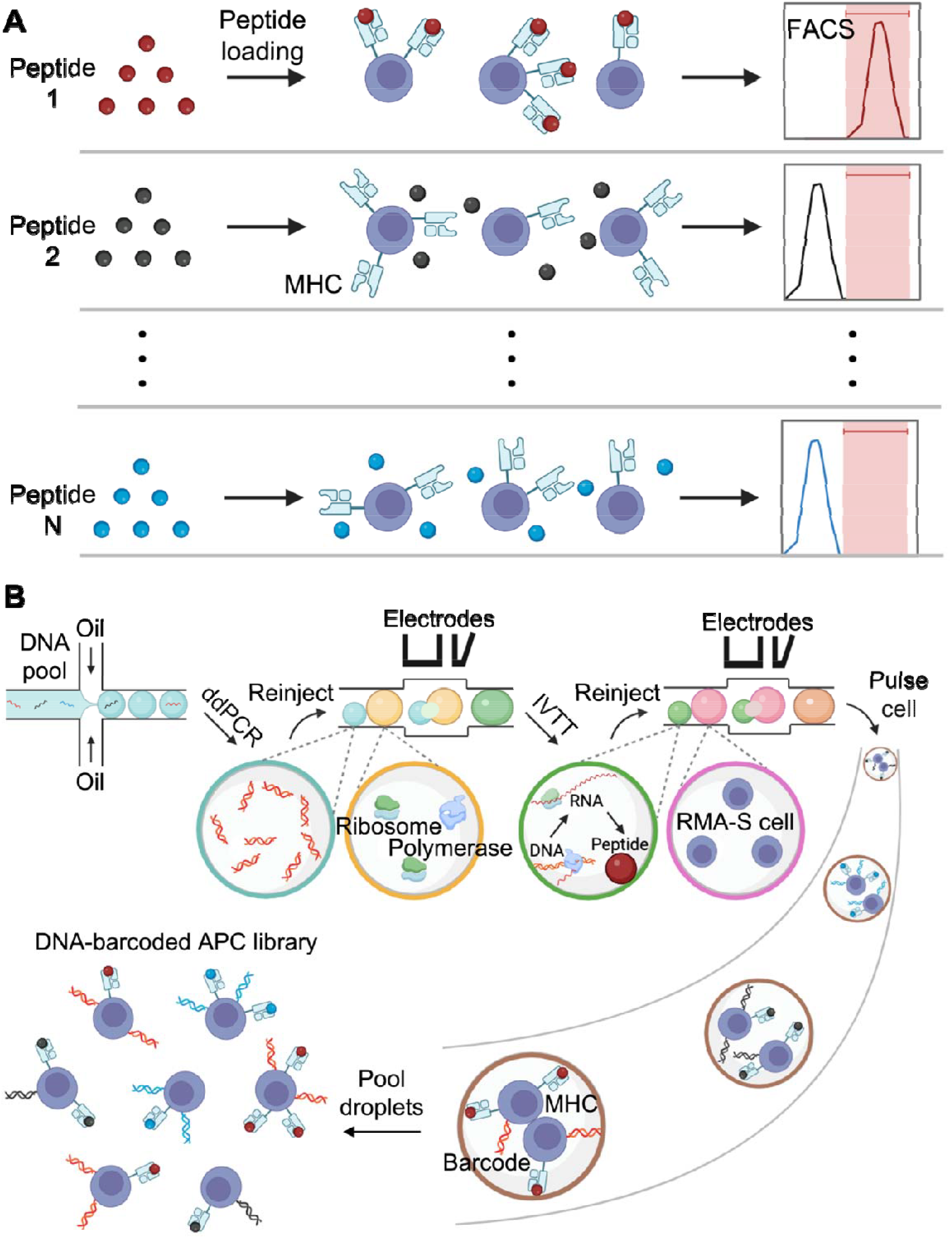
Schematic of the assay. **A)** The conventional approach utilizes FACS to serially analyze APCs pulsed with synthetic peptides. **B)** MEDAL assay makes use of droplet microfluidics to create a library of DNA-barcoded APCs loaded with different peptides. PCR-IVTT provides a rapid and low-cost alternative to produce peptides enzymatically. Simultaneous affinity analysis of multiple peptides is achieved by next-generation sequencing of DNA barcodes in sorted APCs.

We use RMA-S, a TAP-deficient mouse lymphoma cell line expressing the H-2Kb MHC 1 molecule (11) as the APC. Due to its defective antigen-processing mechanism, TAP-deficient cells do not display any endogenous peptides on their surface. Instead, stable MHC-peptide complexes can be formed on the cell surface by external peptide pulsing (12). An important property of MHC molecules is that their stability depends upon the presence of a suitable peptide in its binding cleft. This is determined by the length and amino acid sequence of the peptide fragment, as well as the MHC allele. Without a suitable peptide, the cell-surface MHC molecule will be destabilized and degraded. However, when RMA-S cells are incubated with exogenous peptides capable of binding to the MHC, the peptide-MHC complexes are stabilized and easily detected by flow cytometry with an anti-MHC antibody. Hence, the peptide-induced MHC stability assay is one of the simplest methods to test peptide binding to MHC alleles (13).

We first performed the peptide-induced MHC stability assay on RMA-S cells with chemically synthesized peptides. The mouse H-2Kb restricted OVA peptide (SIINFEKL, ovalbumin 257-264(14)) and human HLA-A2 restricted HBV peptide (FLLTRILTI, Hepatitis B virus 183-191(15)) were chosen as the positive and negative control respectively. RMA-S cells were pulsed with 1 μM synthetic peptides, stained with anti-H-2Kb antibodies, and analyzed with flow cytometry for the peptide-induced MHC stabilization. Our results (Fig 2A) showed that H-2Kb staining in HBV peptide-pulsed RMA-S cells was indistinguishable from control, while a large proportion of RMA-S cells pulsed with OVA peptide showed positive staining, validating the peptide-induced MHC stabilization assay.

**Fig. 2.**
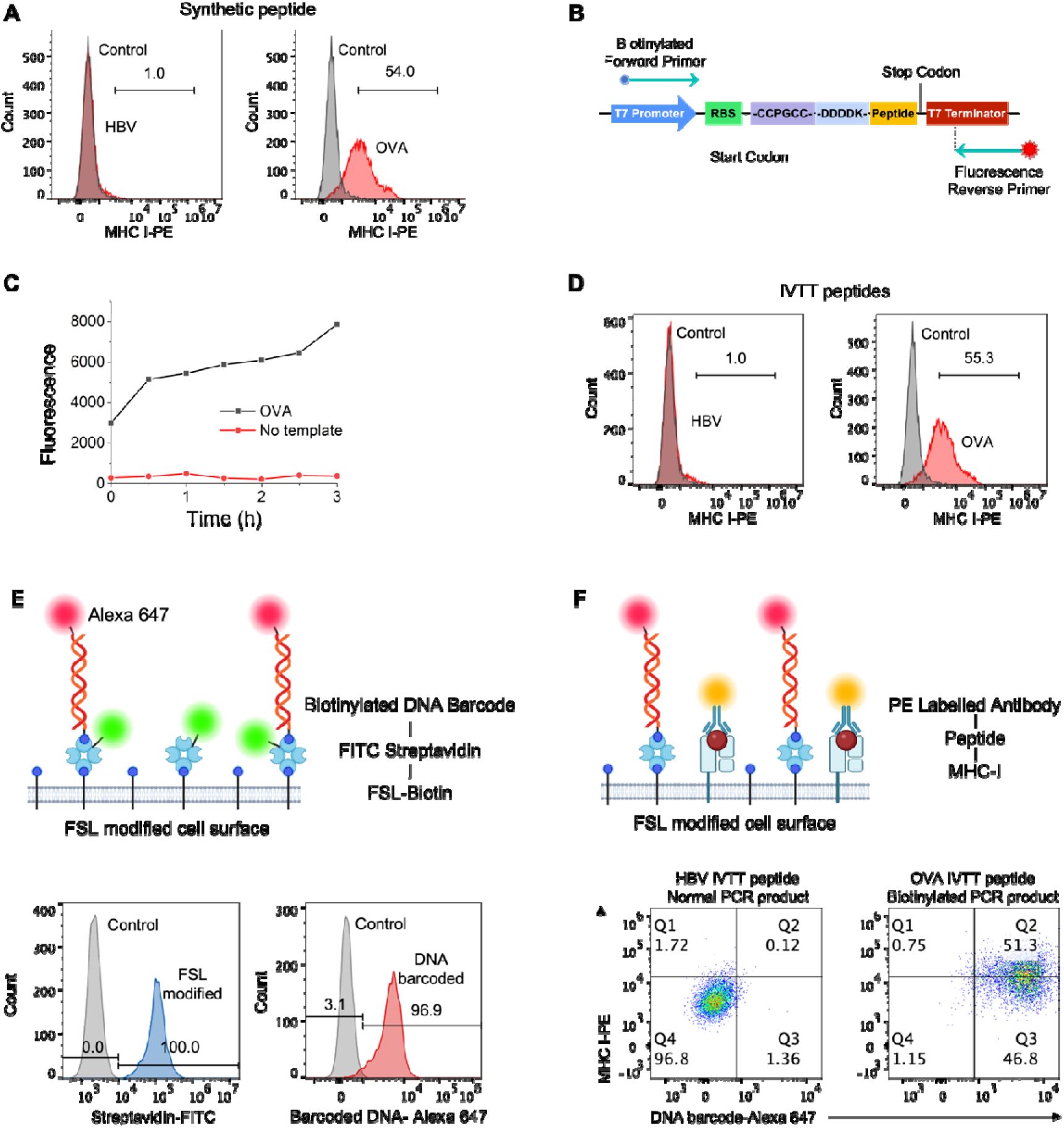
Creating DNA-encoded APC libraries. **A)** Negative HBV peptide binding and positive OVA peptide binding to H-2Kb MHC on RMA-S cells, as demonstrated by peptide-induced MHC-stabilization assay. Chemically synthesized peptides were tested. **B)** Design of DNA template used for PCR-IVTT reaction. **C)** ReAsh signal accumulation during IVTT process from PCR product indicating continuous enzymatic peptide synthesis. **D)** Peptides synthesized from PCR-IVTT reaction are equivalent to chemically synthesized peptides in peptide-induced MHC-stabilization assay. **E)** Schematic for functionalizing cells with DNA barcodes. Histograms show near-complete attachment of streptavidin and barcoded DNA on the cell surface. **F)** Schematic and results for simultaneously measuring DNA barcodes and MHC expression on cell surface.

Our assay seeks to make use of enzyme-synthesized peptides for pulsing APCs to reduce the cost of scaling up. This requires a 3-step procedure of PCR (for generating clonal copies of double-stranded DNA), IVTT, and pulsing cells with the peptides. While previous work (7) has demonstrated that peptides synthesized from IVTT can be successfully loaded into MHC-tetramers, they require multiple purification steps (after PCR, after IVTT) that were fundamentally incompatible with an automated high-throughput workflow. Here, we investigate the feasibility of integrating all 3 steps without purification steps. We first designed the synthetic DNA template as shown in Fig 2B. All DNA sequences used in this study can be found in Table S1 and Fig S1. This template was flanked by the T7 promoter and T7 terminator to facilitate the IVTT process. The open reading frame of the DNA contained the target peptide sequence, preceded by an enterokinase recognition site (-DDDDK-) to facilitate enzymatic digestion of the peptide to the desired length. We also included DNA sequences encoding a tetracysteine tag (-CCPGCC-) upstream of the enterokinase site. The tetracysteine peptide tag could bind to the biarsenical ligand ReAsH to give a red fluorescent signal (16), providing a readout for real-time peptide synthesis in IVTT. In addition, we designed a biotinylated forward primer and a fluorescently labeled reverse primer (Table S2) for amplification of the DNA template in preparation for attaching fluorescent barcodes to the APCs.

We first tested the reagent compatibility in bulk assay. The PCR reaction was first performed, followed by the addition of IVTT reagent supplemented with RNAse inhibitor, enterokinase, and ReAsh dye. During the incubation period, we observed an increase in ReAsh fluorescence, indicating the continuous enzymatic synthesis of the desired peptide sequence (Fig 2C). Next, we incubated the synthetic peptide in combined PCR-IVTT reagent with RMA-S cells, stained the cells with H-2Kb antibody, and performed flow cytometry analysis of MHC stabilization. Our results showed that pulsing RMA-S cells with OVA peptides synthesized with PCR-IVTT reaction resulted in 55.3% H-2Kb-positive cells, while a parallel experiment with HBV peptides had only 1.0% H-2Kb-positive cells (Fig 2D). The excellent agreement of the PCR-IVTT peptide pulsing experiment with the synthetic peptide pulsing experiment validated the feasibility of this scalable, no-purification approach.

Thus far, we have demonstrated a simplified approach for generating peptide-loaded MHCs on APCs through a simple, no-purification PCR-IVTT reaction. However, the MHC-binding potential of each peptide had to be evaluated through separate flow cytometry experiments, limiting the scalability of such an assay. It would be much preferable if APCS with different peptide-loaded MHC could be pooled and analyzed in a single assay. One strategy to enable this is to label each APC with a DNA barcode as it was loaded with a particular peptide. Multiple barcoded APCs corresponding to different peptide-MHC constructs could then be combined, sorted, and analyzed via next-generation sequencing.

We devised a strategy to simultaneously modify the cell surface with the peptide-encoding DNA. For this purpose, we functionalized RMA-S cells with FSL-biotin (17), a biotin-conjugated lipid that could spontaneously insert into the cell membrane creating a biotinylated surface. Streptavidin could be immobilized on the surface of the biotinylated APC in this manner. By including biotinylated forward primers and fluorescently labeled reverse primers in the PCR reaction, biotinylated fluorescent PCR products could be captured on the surface of the streptavidin-functionalized APCs (Fig S2). With this approach, the presence of cell-surface DNA barcode as well as peptide-induced MHC stabilization can be determined via flow cytometry (Fig 2E).

Flow cytometry analysis showed complete modification of cell surface with streptavidin and capture of biotinylated DNA barcodes using this strategy. Meanwhile, a comparable degree of peptide-induced MHC stabilization was observed in these DNA-barcoded cells (Fig 2F). Our results demonstrate the successful implementation of a workflow allowing DNA-barcode-traceable peptide loading on APCs.

We aimed to utilize next-generation sequencing of DNA barcodes to identify peptide sequences that stabilize the MHC complex. For this purpose, we designed primers (Table S2) that could amplify the DNA barcodes that contain peptide information on selected cells (Fig 3A). These primers were flanked by specific adapter sequences to facilitate next-generation sequencing library preparation. To demonstrate this concept, we separately prepared five populations of barcoded cells loaded with different peptides (Fig. 3B, Fig. S3). Among these peptide sequences, we included OVA (SIINFEKL) as a peptide that is expected to bind strongly to H2Kb, and four other negative controls including scrambled OVA sequence (FEKIILSN) (18), HBV (FLLTRILTI) (19), Tryp1(TWHRYHLL) (18) and TUM (KYQAVTTTL) (20). The five APC preparations were pooled, and the cell population that was positive for both DNA barcode and H-2Kb staining was sorted by flow cytometry (Fig 3C).

**Fig. 3:**
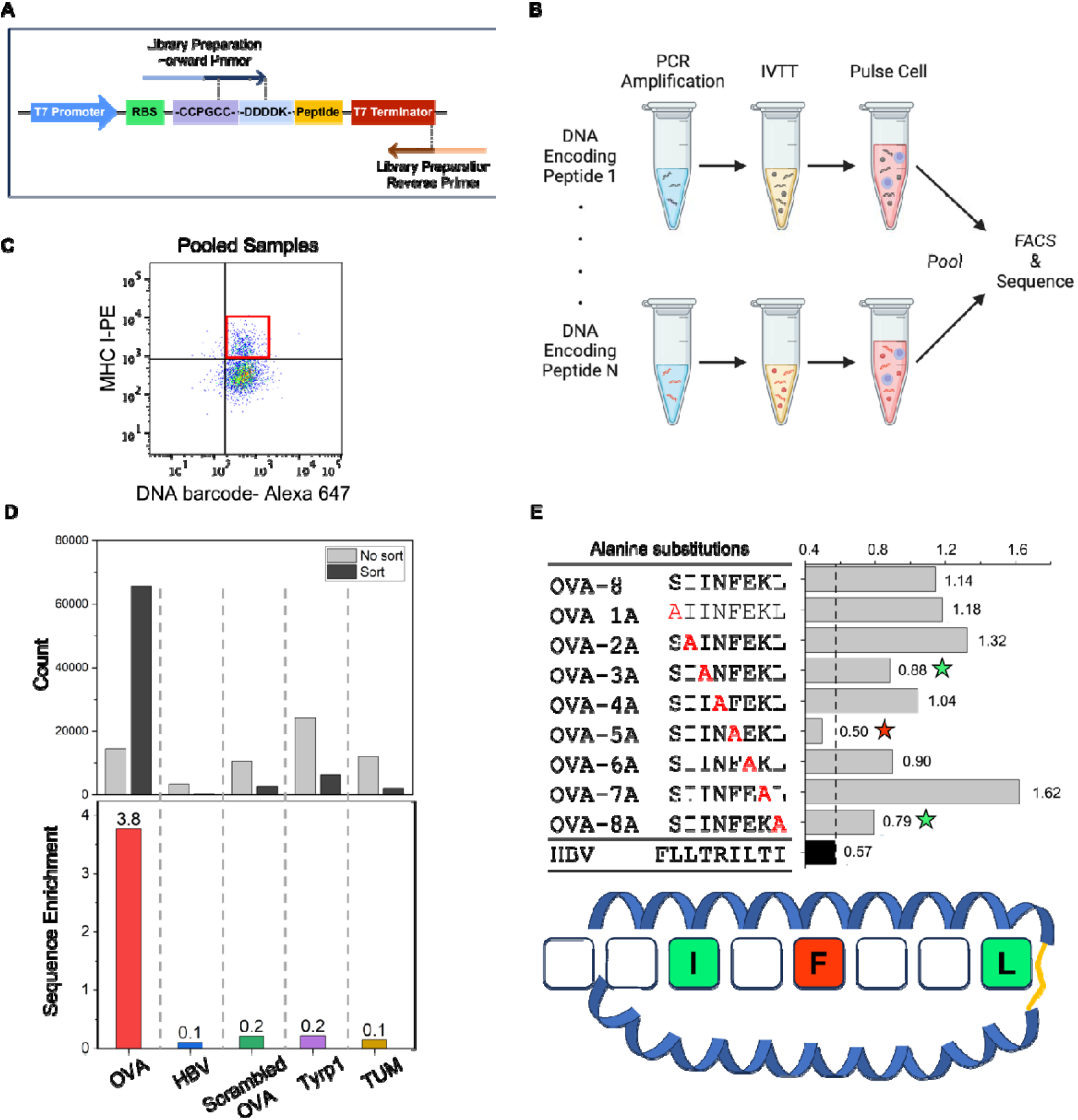
Identifying high-affinity peptides from DNA-barcoded APCs. **A)** Design of primers for amplifying DNA barcodes from cells. B) Multiple pools of DNA-barcoded peptide-loaded APCs are pooled. The population of cells with high MHC expression is sorted by FACS. DNA barcode of the sorted cells is analyzed with next-generation sequencing to identify peptide sequences that bind with high affinity to APCs. **C)** Subpopulation of APCs with high MHC expression in pooled samples. **D)** Sequencing read counts corresponding to the respective peptide sequence in the sorted and unsorted APC pool. OVA peptide shows a high enrichment ratio validating its high affinity to H-2Kb. **E)** Results of Alanine-scanning of OVA peptide on DNA-barcoded APCs. Substitution of F in position 5, L in position 8, and I in position 3 led to a decrease in affinity.

To determine the relative affinity of each peptide to H2Kb, we sequenced the barcodes of both the sorted (enriched for high-affinity peptide) and unsorted cell populations. Fig. 3D showed the uneven distribution of barcode sequences obtained, particularly among the sorted cells. We further defined the enrichment ratio of a particular peptide to be the ratio of normalized reads from the sorted population vs normalized reads from the unsorted population. This yielded a striking enrichment of the OVA peptide sequence compared to the 4 other negative controls. These results demonstrated that our barcode-traceable peptide-loaded APCs could be used for highly multiplexed peptide-MHC affinity assay.

As an alternative application, we also investigated the sensitivity of this platform for determining the contribution of a specific residue in peptide to the stability of the peptide-MHC complex. Using an “Alanine Scanning” approach (21) we generated DNA sequences that encode for peptides that contain substituted amino acid at each position of the 8-residue OVA peptide with Alanine. These 8 DNA sequences encoding alanine-substituted OVA peptide (Table S1), together with a DNA encoding the original OVA peptide and a negative control DNA encoding the HBV peptide sequence were used to prepare barcode-traceable peptide-loaded APCs. Upon flow cytometry analysis of the pooled cells and next-generation sequencing of the sorted and unsorted cell barcodes (Fig S4 and Table S3), we found that substitution of the fifth residue of OVA led to the greatest loss in stability, followed by the third and eighth residues (Fig 3E). Interestingly, this observation was consistent with previous findings, which showed that Phe or Tyr at position 5 and Leu at the carboxy terminus are the two dominant anchor motifs for H2Kb, while position 3 served as a third anchor position (22). This observation provided further evidence of the scalability and sensitivity of our approach.

Our results thus far demonstrated 1) peptide synthesis through IVTT from PCR product, 2) stabilization of MHC on the cell surface by unpurified PCR-IVTT peptide, 3) simultaneous immobilization of barcoded PCR-product on the cell surface, and 4) multiplexed evaluation of peptide-induced MHC stabilization by sequencing the barcode of these cells. Even as these were demonstrated as bulk assays, they were already a significant improvement over other approaches that utilized IVTT for peptide synthesis but required purified PCR product as the template for IVTT and further purification of IVTT reaction for peptide-MHC complexation. Our approach is much more scalable as it does not require purification steps. This advantage is the basis of the MEDAL platform, where the large number of small partitions within a microfluidic platform is exploited for large-scale screening approaches.

To evaluate the feasibility of the MEDAL platform, we developed a three-step procedure consisting of 1) Encapsulating single peptide-encoding DNA from a DNA pool in droplets and performing droplet PCR, 2) Combining droplets containing PCR products with IVTT reagents via droplet fusion to produce the target peptide sequence, and 3) Creating diverse APCs by combining cells with biotinylated PCR product and IVTT-synthesized peptides in individual droplets. APC library was obtained by breaking the emulsions to recover the diverse DNA-barcoded peptide-loaded APCs (Fig 4A). Schematics of all microfluidic devices used in this platform are shown in Fig S5.

**Fig. 4:**
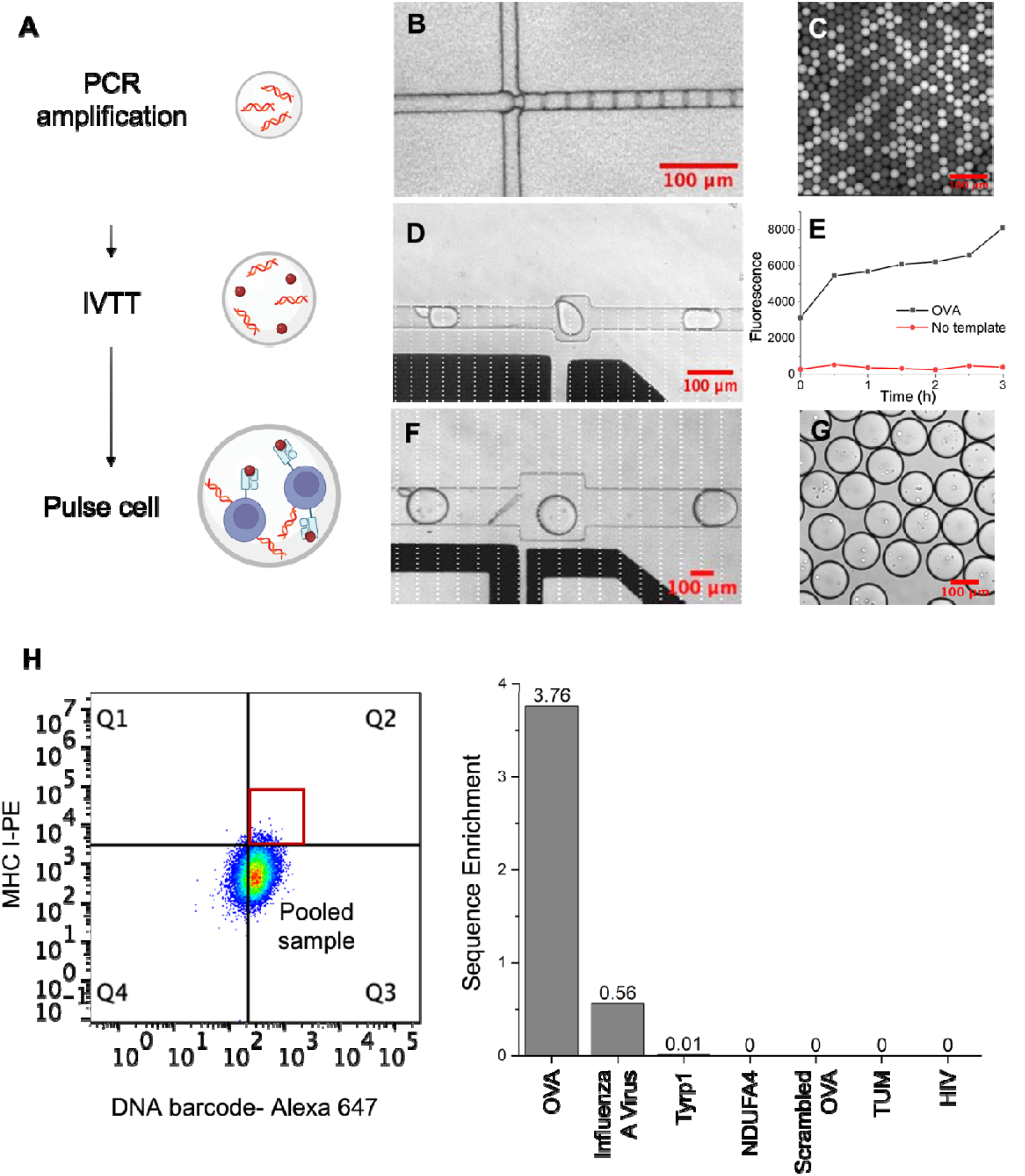
Validation of MEDAL assay. **A)** Schematic of MEDAL assay. **B)** Droplet PCR produces digital amplification of the DNA library in the droplets. **C)** Observe PCR-amplified droplets under the microscope. **D)** Merging of Post-PCR droplets with IVTT droplets under the electric field. **E)** Accumulation of ReAsh dye in PCR-IVTT droplet showing enzymatic peptide synthesis. **F)** Merging of PCR-IVTT droplets with cell droplets under the electric field **G)** Cells encapsulated in PCR-IVTT droplets after merging. **H)** Simultaneous screening of a 7-peptide library using MEDAL. The OVA peptide sequence is highly enriched compared to other peptides.

We performed extensive optimization and validation of each of these steps. Fig. 4B shows the encapsulation of the synthetic DNA template pool in 25 μm droplets. By appropriate adjustment of the template concentration, we ensured that most of the droplets either contained zero or a single initial template, and this was key to generating clonal copies of PCR products in each droplet (Fig 4C). Subsequently, the PCR droplets were collected and injected into a separate chip together with 45 μm diameter droplets containing IVTT reagents. The volume ratio of the PCR and IVTT droplets was designed according to the volume ratio that gave the best peptide synthesis efficiency during our bulk assay optimization. The injection of the PCR and IVTT droplets was synchronized, and electric-field-assisted droplet fusion was carried out in an expanded channel region where both types of droplets were brought to close proximity (23) (Fig 4D, Video S1). We validated that peptide synthesis by IVTT in droplets could be carried out efficiently by monitoring the accumulation of fluorescence signal due to the binding of ReAsh dye to tetracystein tag-containing peptide sequences (Fig 4E).

Finally, APCs were created by combining RMA-S cells (130 μm droplets) with the peptides and DNA in individual droplets (Fig 4F). In this step, peptide-MHC complexation on the cell surface occurred and the cells were concurrently barcoded by the DNA. This process was carried out by an electric-field assisted droplet fusion process similar to the previous step (Video S2 and S3). A high cell concentration (8 million cells/mL) was used to increase the number of barcoded APCs that could be prepared (Fig 4G). After this step, droplet de-emulsification was performed to recover the barcoded APCs. The entire process flow required ∼10 h (2 h for droplet PCR, 4 h for droplet fusion and IVTT, 4 h for droplet fusion and pulsing cells with peptide) and could be used to produce ∼9,000 different kinds of APCs with current parameters.

To validate that this microfluidic platform could produce a properly loaded and barcoded APC pool, we performed this workflow with a pool of DNA templates encoding 7 different peptides (Fig 4H). Among these peptides, OVA is expected to bind and stabilize the H-2Kb MHC, while the rest are predicted to have negligible affinity to the H-2Kb MHC. After recovering the barcoded APCs from droplets, they were stained with anti-H-2Kb antibodies and analyzed with flow cytometry. The cell population that was positive for both DNA barcode and H-2Kb antibody staining was sorted and prepared for DNA barcode sequencing. Next-generation sequencing results showed that compared to the unsorted population, the H-2Kb-positive population was highly enriched for DNA barcodes corresponding to OVA. This peptide-induced MHC stabilization assay demonstrated the successful operation of the MEDAL platform.

## Discussion

Recent success in immunotherapy has created immense interest in harnessing the power of immune cells to treat diseases such as cancer. T cells are particularly attractive as they can recognize immunogenic peptides presented through the cell-surface MHC complex. A current challenge is in linking the arrangement of different amino acid residues in a peptide presented on a cell surface MHC to its immunogenicity. The process is currently laborious and expensive, involving costly individual peptide synthesis for pulsing antigen-presenting cells, and serial functional analysis. Currently, it costs ∼$100 to synthesize a 9-amino acid peptide at 90% purity. The prohibitive cost of conventional inorganic peptide synthesis quickly becomes a bottleneck to scaling these assays.

Enzymatic peptide synthesis, through the use of IVTT reaction, can partially lower this cost. The DNA template used for enzymatic peptide synthesis reagent is ∼10X cheaper than the corresponding synthetic peptide sequence. Using this approach, the production of barcoded MHC-peptide tetramers has been demonstrated (5). Nonetheless, the requirement for multiple reaction purification during the process, coupled with the considerable volumes of expensive reagents needed remains a hurdle to large-scale scaling.

In this manuscript, we present MEDAL, a novel methodology for scalable parallel preparation of barcoded APCs. We demonstrate that a workflow involving template DNA amplification, IVTT peptide synthesis, and cell pulsing with no intermediate purification steps can produce APCs presenting specific peptides. Furthermore, a simple functionalization step enabled the simultaneous engraftment of DNA templates on the cells, creating barcode-traceable APCs. We finally showed that MEDAL can be implemented in a custom-designed microfluidic platform. The capability for parallelization and significant reduction in reaction volume (nL) in a microfluidic platform is key to unprecedented scaling of barcoded APC library preparation.

We have shown a proof of concept of MEDAL for parallel testing of peptide affinity to the H-2Kb MHC among a pool of 7 peptide sequences. The entire workflow can be completed in less than 10 h. Considering the collection of 90,000 droplets and assuming that 10% of the droplets contain DNA templates and synthesized peptides, we estimate that up to 9,000 different APCs can be screened simultaneously.

Currently, a requirement for MEDAL is the use of TAP-deficient cells as APCs to avoid endogenous peptide presentation. The TAP-deficient RMA/S mouse cells expressing the H-2Kb MHC were used here but the equivalent TAP-deficient T2 human cells presenting the HLA-A*0201 MHC are available for human studies (24). Patient-specific Tap-deficient cells (25) can also be produced via CRISPR knockout, opening up the potential to generate personalized barcoded APC libraries.

We envision that the MEDAL platform would also be of significant interest for personalized neoantigen screening. For example, a library of potential neoantigen peptides can be curated based on patient tumor sequencing. A pool of DNA encoding for these peptides can be synthesized at a low cost and used for preparing barcoded APCs as described in this manuscript. Finally, the library of barcoded APCs can be co-incubated with patient T cells. The T cells will kill APCs bearing immunogenic neoantigen peptides, leading to a depletion of the corresponding DNA barcodes. We anticipate that such high-throughput identification of personalized neoantigens would be an important tool for advancing immunotherapy such as in the generation of patient-specific cancer vaccines.

## Experimental Section

### RMA-S cell culture

RMA/S cells were cultured in media (Gibco RPMI Medium 1640 containing 10% Gibco FBS and 1% Gibco Pen Strep) at 37°C. The cells were passaged every five days, and cells with fewer than 10 passages were used for experiments.

### PCR reaction

PCR mastermix contained 0.25 μM of Forward primer and Reverse primer, 1X QX200 ddPCR EvaGreen Supermix, and 0.5 μL template (10 ng/μL). The PCR thermal conditions used were 95 °C for 5 min, 40 cycles of 95°C for 30 s and 60°C for 1min, 4°C for 5 min, 90 °C for 5 min, and hold at 4 °C (Applied Biosystems ProFlex PCR System). The primer sequences can be found in Table 1.

### IVTT reaction

7.3 μL volume of PURExpress IVTT master mix (New England Biolabs) consisted of 4 μL solution A, 3 μL solution B, 0.15 μL RNAse OUT (Invitrogen, Lot no 1908098, 40 U/μL), 0.15 μL Enterokinase (BioLabs, Lot: 10153225, 16000 U/mL). 0.63 μL of Streptavidin FITC Conjugate (Invitrogen, Lot: 2158910) and 2.7 μL of PCR product were added for bulk reactions. Then reaction was incubated at 37°C for 3 h to perform peptide synthesis followed by 23°C incubation for 30 min to facilitate the binding of biotinylated PCR product to FITC Streptavidin.

### RMA-S cell surface functionalization

0.25 million RMA-S cells were washed with 1X PBS (Gibco, PH 7.2) and resuspended in 250 μL 1X PBS solution containing 0.01 mg/mL FSL-biotin (Merck). The cells were incubated at 37°C for 45 min. Next, cells were washed in 1X PBS and resuspended in 250 μL culture media.

### Peptide-induced MHC stabilization assay

To pulse cells with synthetic peptides, synthetic peptides were added to a final concentration of 1 μM to 0.25 million RMA-S cells in culture media. To pulse cells with IVTT peptides, 10 μL of PCR-IVTT reagent was added to 0.25 million RMA-S cells in culture media. The mixture was then incubated at 26°C for 1.5 h and 37°C for 0.5 h.

### Antibody staining and FACS sorting

After the incubation period, cells were pooled together (if multiple samples were used) and subjected to two washes with 1X PBS (Gibco, pH 7.2). Cell quantification was performed using a C-chip (NanoEntek). The volume of added antibody for staining was adjusted based on the tested cell amount, typically using 50 μL of PE anti-mouse H-2Kb (BioLegend, 1 μg/mL) for staining 0.25 million cells on ice for 30 min. Subsequently, cells were washed twice with PBS and resuspended in 450 μL PBS for FACS sorting using the MoFlo Astrios Cell Sorter (Beckman Coulter). Cells were sorted into collection buffer comprising 150 μL of PBS and 20 μL of proteinase K.

### Sequencing library preparation

The collected cells were lysed using the DNeasy Blood & Tissue Kit (Qiagen) following the manufacturer’s instructions. Following that, a next-generation sequencing library was prepared from the extracted DNA. The library preparation PCR master mix included 250 nM of Forward primer and Reverse primer, 1X Gotaq master mix (Promega), and 18 μL of the extracted sample in a final volume of 60 μL. The PCR thermal condition was 95 °C for 2 min, followed by 35 cycles of 95°C for 30 s, 67°C for 30 s, and 72°C for 30 s, with a final extension at 72°C for 5 min. (Applied Biosystems™ ProFlex™ PCR System). The PCR reaction was finally purified with 1.4X SPRI beads (Beckman Coulter) following the provided protocols.

### Droplet assay

#### Device fabrication

Three microfluidic devices were used, including a droplet generator (30 μm height), IVTT merger (30 μm height), and cell merger (45 μm height). The fabrication of these devices employed soft lithography using photomasks (Artnet Pro). The features on the photomask were then transferred to a negative photoresist (Kayaku Advanced Materials, SU-8 2025) on a silicon wafer using UV photolithography (MJB4, SUSS MicroTec).

Polydimethylsiloxane (PDMS, Dow Corning, Sylgard 184) prepolymer mixture was poured over the patterned silicon wafer and cured in a 65 □ °C oven for 2 h. The resulting PDMS replica was peeled off, and inlets and outlets were created by punching with a 1 mm biopsy core. This PDMS piece was bonded to a PDMS-coated glass slide using an oxygen plasma cleaner (PDC-32G, Harrick), followed by baking at 65 □ °C for 1 h to ensure robust bonding.

For the electrode components, indium alloy wire (Indium Corporation) was melted into the electrode inlet channels and connected to wires.

#### Droplet digital PCR

25 μM diameter droplets were generated by using QX200™ Droplet Generation Oil (used in all microfluidic devices) for EvaGreen. The flow rates employed were 48 μL/h for the aqueous phase and 480 μL/h for the oil phase. To ensure a Poisson distribution, a DNA template concentration of 1 pg/μL was added to the PCR master mix, following the same thermal conditions as those used in the bulk assay.

#### Merging PCR with IVTT droplets

45 μM diameter droplets containing IVTT master mix were produced by setting flow rates of 60 μL/h for the aqueous phase (IVTT master mix) and 240 μL/h for the oil phase. PCR droplets were reinjected into the device at a flow rate of 12 μL/h and oil was injected at a flow rate of 60 μL/h to space the reinjected droplets. IVTT droplets and PCR droplets were merged at the designated location within the chip with a 15 VPP 1.5 kHz sinusoidal electric field applied through the electrodes. Droplets were collected from the chip outlet and incubated at 37°C for 3 h and 23°C for 30 min.

#### Merging PCR-IVTT with cell droplets

130 μM diameter droplets containing RMA-S cells (in RPMI Medium with 16% (v/v) Optiprep (Sigma-Aldrich, St. Louis, MO)) were produced by setting flow rates of 60 μL/h for aqueous phase and 480 μL/h for the oil phase. PCR-IVTT droplets were reinjected into the device at a flow rate of 12 μL/h and oil was injected at a flow rate of 60 μL/h to space the reinjected droplets. Cell-containing droplets and PCR-IVTT droplets were merged at the designated location within the chip with a 10 VPP 1.5 kHz sinusoidal electric field applied through the electrodes. Droplets were collected from the chip outlet and incubated at 26°C for 1.5 h and 37°C for 0.5 h. Subsequently, droplets were de-emulsified in 1H,1H,2H,2H-Perfluoro-1-octanol reagent (PFO, Sigma-Aldrich). Collected cells were stained with PE anti-mouse H-2Kb, BioLegend, 1 μg/mL) for FACS sorting.

#### Monitoring peptide synthesis in droplets with ReAsh

Droplets containing PCR product (or no template controls) were merged with droplets containing IVTT master mix according to the established protocol for the droplet assay. ReAsh-EDT2 (Thermo Fisher Scientific) was added to the IVTT mix to achieve a final concentration of 10 μM. The resulting droplets were incubated at 37°C for varying durations, de-emulsified with PFO treatment, and fluorescence intensity was quantified using a TECAN plate reader.

## Acknowledgments

This work was supported by funding from the Ministry of Education of Singapore (MOE-T2EP30120-0008)

## References

1. A.-L. Schaap-Johansen, M. Vujović, A. Borch, S. R. Hadrup, P. Marcatili, T Cell Epitope Prediction and Its Application to Immunotherapy. Front Immunol 12 (2021).

2. D. Hu, A. T. Irving, Massively-multiplexed epitope mapping techniques for viral antigen discovery. Front Immunol 14 (2023).

3. E. W. Newell, L. O. Klein, W. Yu, M. M. Davis, Simultaneous detection of many T-cell specificities using combinatorial tetramer staining. Nat Methods 6, 497–499 (2009).

4. E. W. Newell, N. Sigal, N. Nair, B. A. Kidd, H. B. Greenberg, M. M. Davis, Combinatorial tetramer staining and mass cytometry analysis facilitate T-cell epitope mapping and characterization. Nat Biotechnol 31, 623–629 (2013).

5. A. K. Bentzen, A. M. Marquard, R. Lyngaa, S. K. Saini, S. Ramskov, M. Donia, L. Such, A. J. S. Furness, N. McGranahan, R. Rosenthal, P. thor Straten, Z. Szallasi, I. M. Svane, C. Swanton, S. A. Quezada, S. N. Jakobsen, A. C. Eklund, S. R. Hadrup, Large-scale detection of antigen-specific T cells using peptide-MHC-I multimers labeled with DNA barcodes. Nat Biotechnol 34, 1037–1045 (2016).

6. B. Rodenko, M. Toebes, S. R. Hadrup, W. J. E. van Esch, A. M. Molenaar, T. N. M. Schumacher, H. Ovaa, Generation of peptide–MHC class I complexes through UV-mediated ligand exchange. Nat Protoc 1, 1120–1132 (2006).

7. S.-Q. Zhang, K.-Y. Ma, A. A. Schonnesen, M. Zhang, C. He, E. Sun, C. M. Williams, W. Jia, N. Jiang, High-throughput determination of the antigen specificities of T cell receptors in single cells. Nat Biotechnol 36, 1156–1159 (2018).

8. Y. Xu, K.-C. Wu, W. Jiang, Y. Hou, L. F. Cheow, V. H.-F. Lee, C.-H. Chen, Single-Cell Secretion Analysis via Microfluidic Cell Membrane Immunosorbent Assay for Immune Profiling. Anal Chem 96, 49–58 (2024).

9. W. N. Lin, M. Z. Tay, J. X. E. Wong, C. Y. Lee, S.-W. Fong, C.-I. Wang, L. F. P. Ng, L. Renia, C.-H. Chen, L. F. Cheow, Rapid microfluidic platform for screening and enrichment of cells secreting virus neutralizing antibodies. Lab Chip 22, 2578–2589 (2022).

10. B. Laugel, H. A. van den Berg, E. Gostick, D. K. Cole, L. Wooldridge, J. Boulter, A. Milicic, D. A. Price, A. K. Sewell, Different T Cell Receptor Affinity Thresholds and CD8 Coreceptor Dependence Govern Cytotoxic T Lymphocyte Activation and Tetramer Binding Properties. Journal of Biological Chemistry 282, 23799–23810 (2007).

11. H.-G. Ljunggren, C. Öhlén, P. Höglund, L. Franksson, K. Kärre, The RMA-S lymphoma mutant; consequences of a peptide loading defect on immunological recognition and graft rejection. Int J Cancer 47, 38–44 (1991).

12. T. N. M. Schumacher, M.-T. Heemels, J. J. Neefjes, W. M. Kast, C. J. M. Melief, H. L. Ploegh, Direct binding of peptide to empty MHC class I molecules on intact cells and in vitro. Cell 62, 563–567 (1990).

13. M. Harndahl, M. Rasmussen, G. Roder, I. Dalgaard Pedersen, M. Sørensen, M. Nielsen, S. Buus, PeptideLMHC class I stability is a better predictor than peptide affinity of CTL immunogenicity. Eur J Immunol 42, 1405–1416 (2012).

14. O. Rötzschke, K. Falk, S. Stevanovic, G. Jung, P. Walden, H. Rammensee, Exact prediction of a natural T cell epitope. Eur J Immunol 21, 2891–2894 (1991).

15. R. Bertoni, J. Sidney, P. Fowler, R. W. Chesnut, F. V Chisari, A. Sette, Human histocompatibility leukocyte antigen-binding supermotifs predict broadly cross-reactive cytotoxic T lymphocyte responses in patients with acute hepatitis. Journal of Clinical Investigation 100, 503–513 (1997).

16. S. R. Adams, R. E. Campbell, L. A. Gross, B. R. Martin, G. K. Walkup, Y. Yao, J. Llopis, R. Y. Tsien, New Biarsenical Ligands and Tetracysteine Motifs for Protein Labeling in Vitro and in Vivo: Synthesis and Biological Applications. J Am Chem Soc 124, 6063–6076 (2002).

17. D. A. Blake, N. V. Bovin, D. Bess, S. M. Henry, FSL Constructs: A Simple Method for Modifying Cell/Virion Surfaces with a Range of Biological Markers Without Affecting their Viability. Journal of Visualized Experiments, doi: 10.3791/3289 (2011).

18. R. S. Gejman, A. Y. Chang, H. F. Jones, K. DiKun, A. A. Hakimi, A. Schietinger, D. A. Scheinberg, Rejection of immunogenic tumor clones is limited by clonal fraction. Elife 7 (2018).

19. R. Schirmbeck, J. Reimann, S. Kochanek, F. Kreppel, The Immunogenicity of Adenovirus Vectors Limits the Multispecificity of CD8 T-cell Responses to Vector-encoded Transgenic Antigens. Molecular Therapy 16, 1609–1616 (2008).

20. Y. Saito, P. A. Peterson, M. Matsumura, Quantitation of peptide anchor residue contributions to class I major histocompatibility complex molecule binding. J Biol Chem 268, 21309–17 (1993).

21. B. C. Cunningham, J. A. Wells, High-Resolution Epitope Mapping of hGH-Receptor Interactions by Alanine-Scanning Mutagenesis. Science (1979) 244, 1081–1085 (1989).

22. S. C. Jameson, M. J. Bevan, Dissection of major histocompatibility complex (MHC) and T cell receptor contact residues in a K b Lrestricted ovalbumin peptide and an assessment of the predictive power of MHCLbinding motifs. Eur J Immunol 22, 2663–2667 (1992).

23. E. Brouzes, M. Medkova, N. Savenelli, D. Marran, M. Twardowski, J. B. Hutchison, J. M. Rothberg, D. R. Link, N. Perrimon, M. L. Samuels, Droplet microfluidic technology for single-cell high-throughput screening. Proceedings of the National Academy of Sciences 106, 14195–14200 (2009).

24. R. A. Henderson, H. Michel, K. Sakaguchi, J. Shabanowitz, E. Appella, D. F. Hunt, V. H. Engelhard, HLA-A2.1-Associated Peptides from a Mutant Cell Line: A Second Pathway of Antigen Presentation. Science (1979) 255, 1264–1266 (1992).

25. C. Kaseke, R. J. Park, N. K. Singh, D. Koundakjian, A. Bashirova, W. F. Garcia Beltran, O. C. Takou Mbah, J. Ma, F. Senjobe, J. M. Urbach, A. Nathan, E. J. Rossin, R. Tano-Menka, A. Khatri, A. Piechocka-Trocha, M. T. Waring, M. E. Birnbaum, B. M. Baker, M. Carrington, B. D. Walker, G. D. Gaiha, HLA class-I-peptide stability mediates CD8+ T cell immunodominance hierarchies and facilitates HLA-associated immune control of HIV. Cell Rep 36, 109378 (2021).

