## Supplementary Materials for "Microfluidic-enabled production of DNA barcoded APC library (MEDAL) for high throughput T cell epitope screening"

Xu Cui *et al.*

**This PDF file includes:**

Table S1 to S4

Fig S1 to S5

Captions for Video S1 to S3

**Other Supplementary Materials for this manuscript include the following:**

Video S1 to S3

**Table S1.**

DNA sequences encoding for selected peptides.

| Peptide name | Peptide sequence | Template sequence |
| --- | --- | --- |
| OVA | SIINFEKL | TCCATCATTAACTTTGAGAAGCTG |
| OVA-1A | AIINFEKL | GCGATCATTAACTTTGAGAAGCTG |
| OVA-2A | SAINFEKL | TCCGCGATTAACTTTGAGAAGCTG |
| OVA-3A | SIANFEKL | TCC ATCGCGAACTTTGAGAAGCTG |
| OVA-4A | SIIAFEKL | TCCATCATTGCGTTTGAGAAGCTG |
| OVA-5A | SIINAEKL | TCCATCATTAACGCGGAGAAGCTG |
| OVA-6A | SIINFAKL | TCCATCATTAACTTTGCGAAGCTG |
| OVA-7A | SIINFEAL | TCCATCATTAACTTTGAGGCGCTG |
| OVA-8A | SIINFEKA | TCCATCATTAACTTTGAGAAGGCG |
| HBV | FLLTRILTI | TTTCTGCTGACCCGCATTCTGACCATT |
| TUM | KYQAVTTTL | AAATATCAGGCGGTGACCACCACCCTG |
| Influenza A Virus | ASNENMETM | GCGAGCAACGAAAACATGGAAACCATG |
| HIV | ILKEPVHGV | ATTCTGAAAGAACCGGTGCATGGCGTG |
| Scrambled OVA | FEKIILSN | TTTGAAAAAATTATTCTGAGCAAC |
| NDUFA4 | EQYKFYSV | GAACAGTATAAATTTTATAGCGTG |
| Tyrp 1 | TWHRYHLL | ACCTGGCATCGCTATCATCTGCTG |

**Table S2.**

Primer sequences used in this study.

| **Primers for DNA templates amplification** | **Oligo sequence** |
| --- | --- |
| Biotinylated forward primer | 5-' (/5Biosg/) GCG AAA TTA ATA CGA CTC ACT ATA GG -3' |
| Fluorescent reverse primer | 5-' /5Alex647N/CAA AAA ACC CCTCAA GAC C -3' |

| **Primers for library preparation** | **Oligo sequence** |
| --- | --- |
| Library preparation forward primer | AATGATACGGCGACCACCGAGATCTACACTCTTTCCCTACACGACGCTCTTCCGATCTGTTGCTGTGATGATGATGAT |
| Library preparation reverse primer | CAAGCAGAAGACGGCATACGAGAT[6 bp Index]GTGA  CTGGAGTTCAGACGTGTGCTCTTCCGATCT CAAAAAACCCCTCAAGACCC |

**Table S3.**

Sequencing read counts of DNA corresponding to peptides in alanine-substitution library.

| Sequence | OVA-8 | OVA-1A | OVA-2A | OVA-3A | OVA-4A | OVA-5A | OVA-6A | OVA-7A | OVA-8A | HBV |
| --- | --- | --- | --- | --- | --- | --- | --- | --- | --- | --- |
| Sorted cells | 38651 | 47720 | 75783 | 46420 | 42183 | 20287 | 29552 | 63022 | 24446 | 26954 |
| Unsorted cells | 47635 | 56957 | 80678 | 74180 | 57086 | 57701 | 46437 | 54835 | 43666 | 67055 |

**Table S4.**

Sequencing read counts of DNA corresponding to peptides in droplet MEDAL assay.

| Sequence | OVA | Influenza A Virus | Tyrp 1 | NDUFA4 | Scrambled OVA | TUM | HIV |
| --- | --- | --- | --- | --- | --- | --- | --- |
| Sorted cells | 32620 | 5230 | 145 | 0 | 0 | 0 | 0 |
| Unsorted cells | 8931 | 9577 | 12930 | 1587 | 233 | 2297 | 3595 |

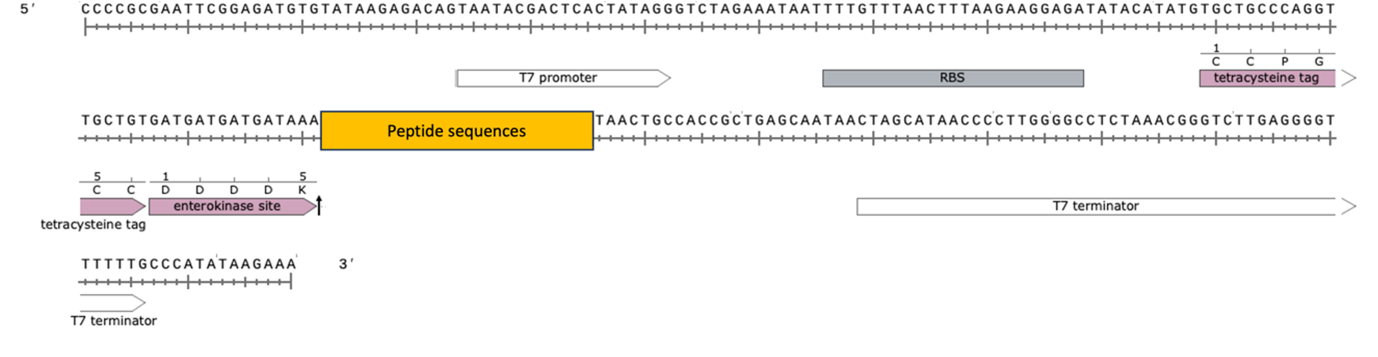

Fig S1. Sequence and design of IVTT-compatible DNA template encoding peptides.

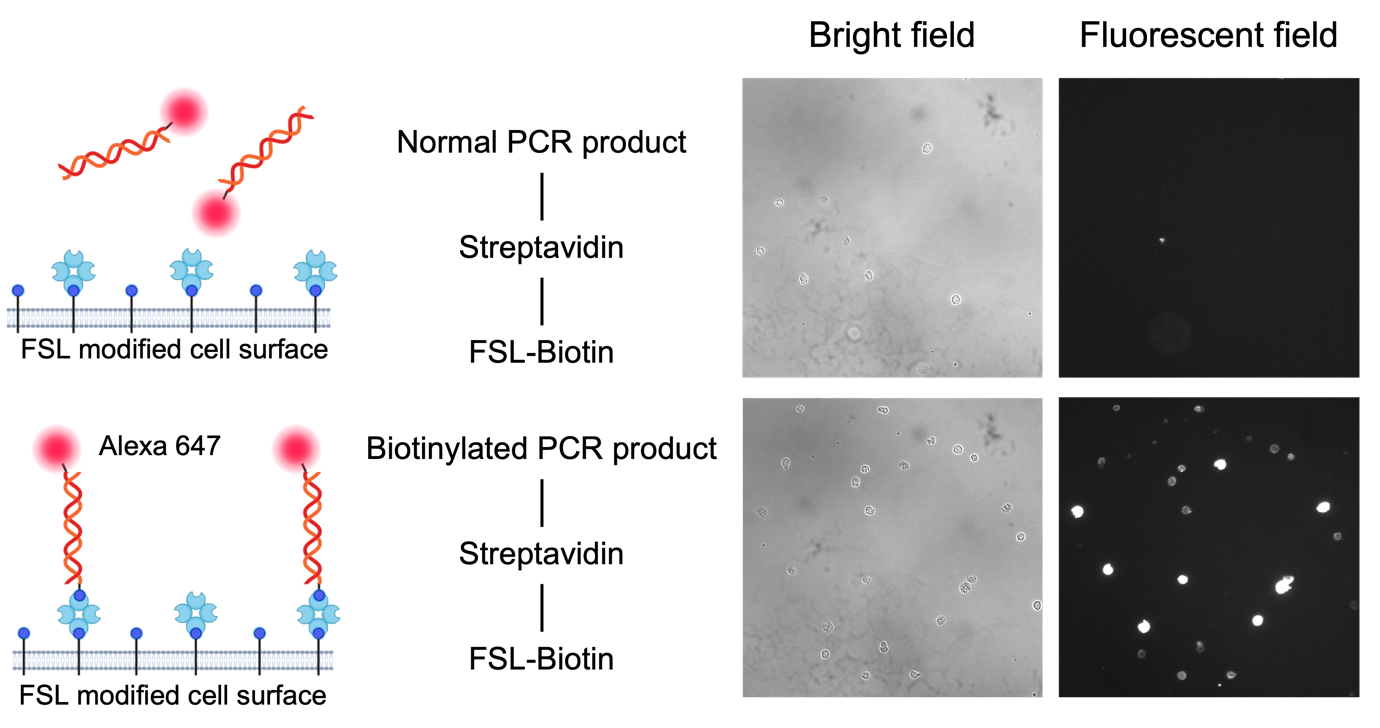

**Fig S2.** **Pre-functionalization of cells to allow attachment of DNA barcodes.** RMA-S cells modified with FSL-biotin was functionalized with streptavidin, and further incubated with fluorescent PCR products with or without biotin modification. Only the use of biotin-modified PCR product led to their attachment to cell surface.

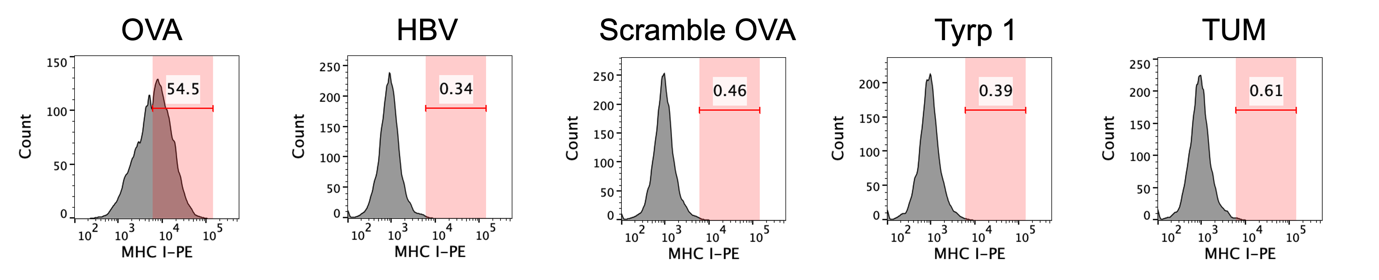

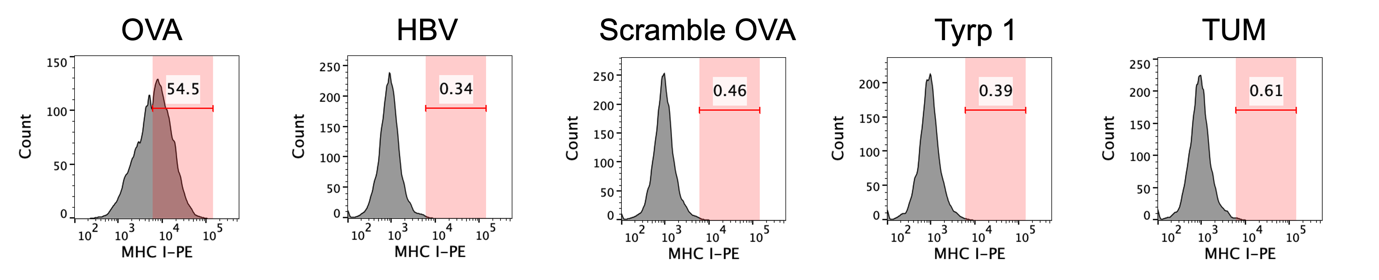

**Fig S3.** Histogram of H-2Kb staining in RMA-S cells pulsed with different kinds of IVTT peptides. The binding of high-affinity OVA peptide to H-2Kb on the cell surface led to their stabilization compared to other peptides.

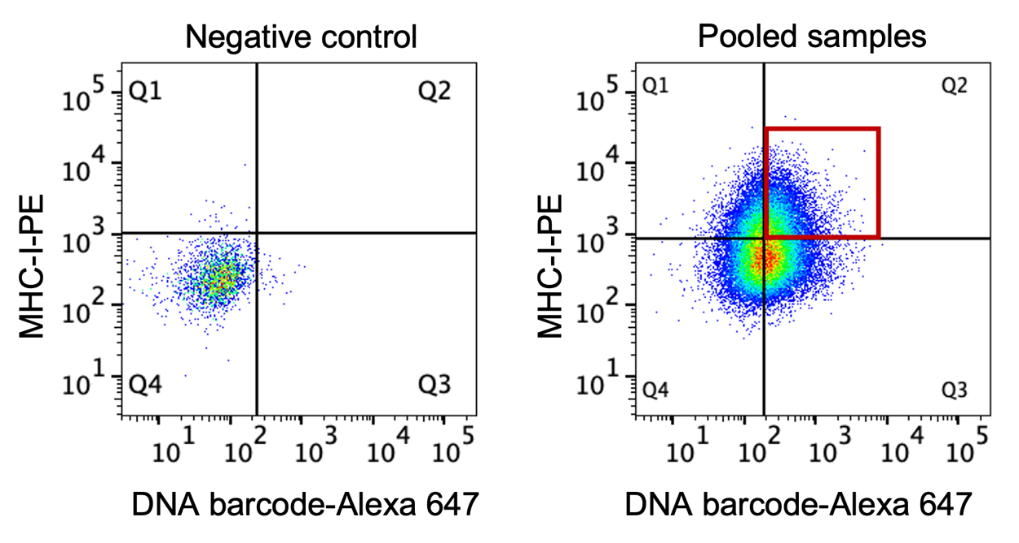

**Fig S4. Gating strategy for sorting cells with high H-2Kb expression.** Left: H-2Kb and DNA barcode signals in unmodified cells. Right: H-2Kb and DNA barcode signals in the pool of cells incubated with the alanine-substitution library and its corresponding DNA barcode. The red box indicates cells that were FACS sorted for DNA sequencing.

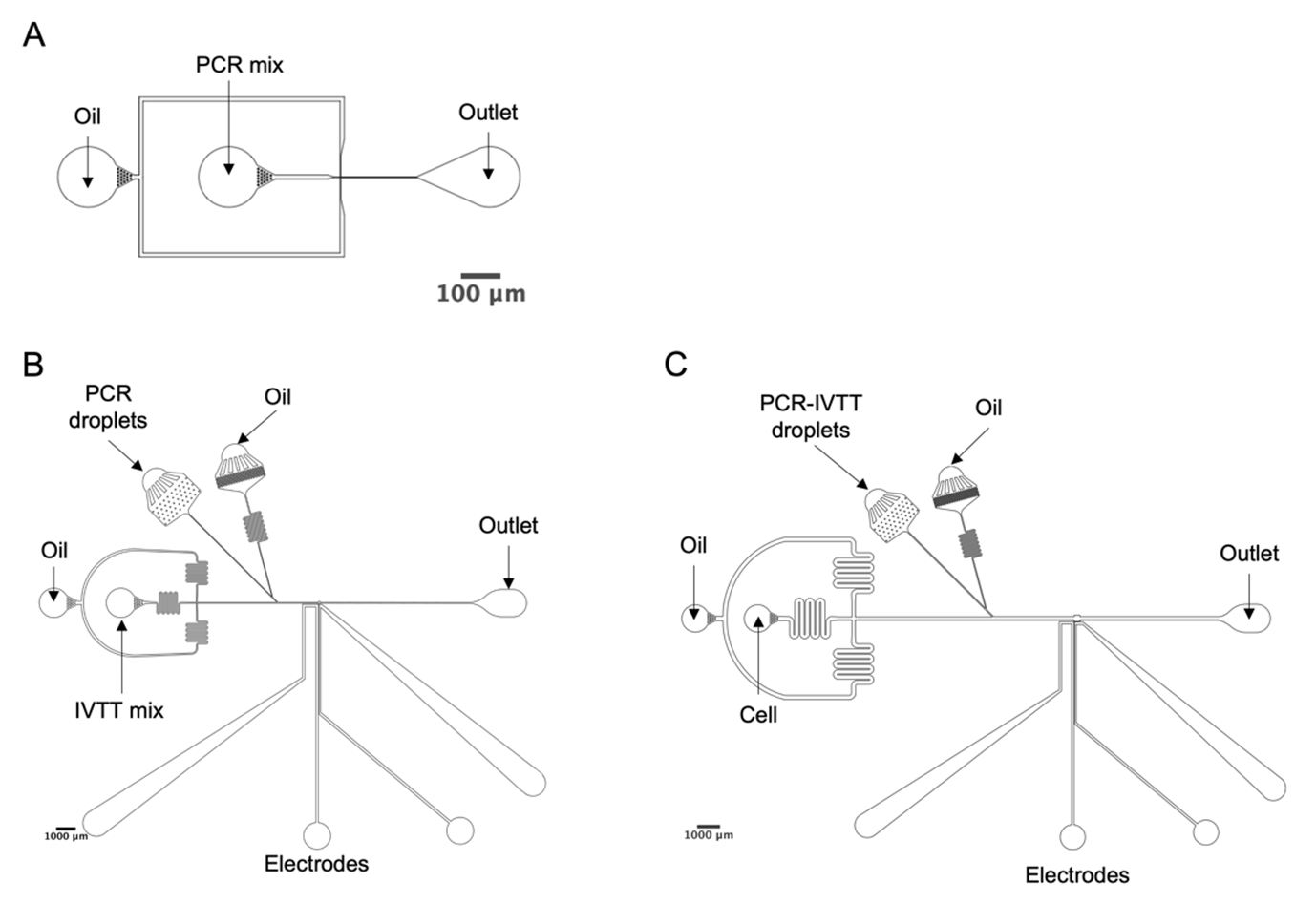

**Fig S5.** Schematics of microfluidic chips: A. droplet generator, B. IVTT merger, and C. Cell merger.
